# *TET1* and *TDG* suppress intestinal tumorigenesis by down-regulating the inflammatory and immune response pathways

**DOI:** 10.1101/676445

**Authors:** Rossella Tricarico, Jozef Madzo, Gabrielle Scher, Shinji Maegawa, Jaroslav Jelinek, Carly Scher, Wen-Chi Chang, Emmanuelle Nicolas, Yan Zhou, Michael Slifker, Karthik Devarajan, Kathy Q. Cai, Pamela Nakajima, Jinfei Xu, Pietro Mancuso, Valentina Doneddu, Luigi Bagella, Justin Ingram, Siddharth Balachandran, Iuliia Peshkova, Ekaterina Koltsova, Sergei Grivennikov, Timothy J. Yen, Jean-Pierre Issa, Alfonso Bellacosa

## Abstract

**Introduction:** Aberrant DNA methylation is frequently observed in colorectal cancer (CRC), but the underlying mechanisms are poorly understood. Ten-Eleven Translocation (TET) dioxygenases and DNA repair enzyme Thymine DNA Glycosylase (TDG) are involved in active DNA demethylation by generating and removing, respectively, novel oxidized cytosine species. Mutations of *TET1* and *TDG*, and alterations of the levels of oxidized cytosine species have been identified in human CRC cases, but the biological significance of the TET-TDG demethylation axis in intestinal tumorigenesis is unclear.

**Material and Methods:** We generated *Apc*^Min^ mice with additional inactivation of *Tet1* and/or *Tdg*, and characterized the methylome and transcriptome of intestinal adenomas by DREAM and RNA sequencing, respectively.

**Results:** *Tet1-* and/or *Tdg*-deficient *Apc*^Min^ mice show enhanced intestinal tumorigenesis in comparison to wild type *Tet1* and *Tdg Apc*^Min^ mice. Specifically, *Tet1* and/or *Tdg*-deficient *Apc*^Min^ adenomas manifested increased size or features of erosion and stroma activation. Methylome analysis revealed progressive loss of global DNA hypomethylation in colonic adenomas from *Tet1-* and *Tdg*-deficient *Apc*^Min^ mice, and hypermethylation of CpG islands in *Tet1*-deficient *Apc*^Min^ mice. In addition, RNA sequencing showed upregulation of genes in inflammatory and immune response pathways in *Tet1*-and *Tdg*-mutant colonic adenomas compared to control *Apc*^Min^ adenomas.

**Conclusions:** Taken together, these findings demonstrate the important role of active DNA demethylation mediated by TET-TDG in reducing intestinal tumor formation, by modulating the epigenome and inflammatory/immune responses. This study highlights a novel mechanism of epigenetic deregulation during intestinal tumorigenesis with diagnostic, therapeutic and prognostic implications.

## Introduction

Colorectal cancer is the second leading cause of death from cancer in Western countries. Every year, approximately 140,000 new cases are diagnosed in U.S.A. and approximately 50,000 deaths occur (1). It is reasonable to assume that successful control of this disease, via appropriate preventive, interventional and therapeutic strategies, will depend on the precise characterization of its molecular basis. Converging studies from different disciplines revealed that colorectal cancer, like many other epithelial malignancies, is characterized by multi-step carcinogenesis. In particular, studies by Drs. Fearon and Vogelstein and Dr. Feinberg and colleagues demonstrated that several genetic and epigenetic alterations accumulate in cancer cells at critical genes that control rate-limiting steps of cell proliferation, differentiation and death (2,3). This has led to the hypothesis that some discrete genetic and epigenetic changes accompany the morphological transitions from normal colonic cells to a benign tumor (adenoma), to a malignant cancer (carcinoma).

Among the known genetic alterations in colorectal cancer development are inactivating mutations of the tumor suppressor genes *APC, SMAD2-4* and *TP53*, and activating mutations of the oncogene *KRAS*. In addition, several epigenetic changes have been described, involving DNA methylation, chromatin modification and remodeling, and microRNA expression. Focusing our attention only on DNA-based epigenetic alterations, we can recognize: genome-wide DNA hypomethylation, particularly evident at satellite sequences; DNA hypermethylation and silencing of promoters of tumor suppressor genes and other CpG-rich sequences; and loss of imprinting marks discriminating the paternal and maternal alleles. Simultaneous hypermethylation of multiple CpG islands, the so called CpG island methylator phenotype (CIMP), is particularly evident in a fraction of colorectal tumors (4). While some hypermethylation events are linked to aging (5), it is clear that even after correcting for age, consistent hypermethylation of clusters of CpG islands is found in a subset of tumors. In fact, the existence of CIMP is probably the best evidence of epigenetic instability occurring in tumors (4). Gene silencing by CIMP represents a powerful pathogenetic mechanisms because it can achieve inactivation of multiple tumor suppressor genes at once. CIMP has been recently described by the TCGA in other tumors, including gliomas and breast cancers. The molecular basis of CIMP in CRC is unknown. It is possible that CIMP reflects environmental or dietary exposure to epimutagens. It is also possible that CIMP originates from an imbalance of methylating activities and demethylating activities that normally protect CpG islands from methylation (4).

Ten-Eleven Translocation (TET) family dioxygenases play a crucial role in active DNA demethylation by oxidizing 5-methylcytosine to 5-hydroxymethylcytosine (5hmC) and converting 5hmC to 5-formylcytosine (5fC) and 5-carboxylcytosine (5caC)(6-10). Our data indicate that, through its CXXC domain, TET1 binds preferentially to hypomethylated CpG islands and maintains their hypomethylated status by removing and preventing the spreading of aberrant DNA methylation (11). Hypothetically, loss of TET1 function might lead to the hypermethylator phenotype seen in CIMP. Low levels of 5hmC suggest an involvement of TET enzymes in CRC tumorigenesis (12,13). In addition, we and others recently identified the base excision repair enzyme TDG as a critical effector of DNA demethylation downstream of TET enzymes, through the removal of 5fC, 5caC and 5-hydroxymethyluracil (5hmU), a deamination product of 5hmC (14-19). We also found that the expression of TDG is frequently reduced in CRC cell lines, and that *TDG* acts as an important tumor suppressor in the pathogenesis of a subset of intestinal tumors (20). In addition, inactivation of TDG is associated with increased methylation (21). Thus, loss of TDG could also result in CIMP or other epigenomic changes in CRC tumorigenesis.

Here, to functionally study the role of TET1 and TDG in epigenome modulation and CRC formation, we crossed mice bearing the *Tet1*^*-*^ knock-out allele and/or the dominant-negative, glycosylase-dead *Tdg*^*N151A*^ allele with Apc-Min (Multiple Intestinal Neoplasia) mice, predisposed to colorectal neoplasia (in particular, we used the Fox Chase Cancer Center variant, characterized by high incidence of colonic lesions)(22,23). The results revealed a complex role of TET1 and TDG in modulating methylation patterns and gene expression, and revealed a tumor suppressive role via inhibition of inflammatory and immune response pathways.

## Materials and Methods

### Experimental animals

To generate F_1_ double heterozygotes (*Tet1*^+/-^ *Apc*^Min/+^ and *Tet1*^+/-^ *Tdg*^N151A/+^) mice, *Tet1*^+/-^ (24)(from The Jackson Laboratory) and *Apc*^Min-FCCC^ (22), were mated with *Tdg*^N151A/+^ mice (16). F_1_ *Tet1*^+/-^ *Apc*^Min/+^ and *Tet1*^+/-^ *Tdg*^N151A/+^ were crossed to generate 90 F_2_ mice divided into the following four-genotype groups: *Tet1*^-/-^ *Tdg*^+/+^ *Apc*^Min/+^ (n=7); *Tet1*^+/-^ *Tdg*^N151A/+^ *Apc*^Min/+^ (n=21); *Tet1*^+/+^ *Tdg*^N151A/+^ *Apc*^Min/+^ (n=33); and *Tet1*^+/+^ *Tdg*^+/+^ *Apc*^Min/+^ (n=29). All the strains used in this study have a C57BL/6 genetic background. The Institutional Animal Care and Use Committee of the Fox Chase Cancer Center approved animal protocols and all the procedures of mouse handling.

### Histopathology and tumor load analyses

All the animals were monitored weekly for rectal prolapse and/or bleeding and sacrificed at 150-160 days. At the time of sacrifice, the small intestine, cecum, colorectum and other various tissues (including kidney, liver and spleen) were collected and inspected. Proximal, middle, and distal small intestine and colorectum were embedded in paraffin, and sections were stained with hematoxylin/eosin for histopathological analysis according to standard procedures. Small intestinal and colorectal adenomas were counted, evaluated for hemorrhagic features and measured with a ruler. In addition, colonic adenomas were also collected for molecular analysis and histopathology. A detailed analysis of the intestinal adenomas using standard histopathologic criteria was also performed by Dr. H.S. Cooper, an expert pathologist of intestinal tumors in *Apc*^Min-FCCC^.

### Genome-wide methylation analysis

Genomic DNA was extracted from colonic adenomas and normal mucosa by Gentra Puregene Tissue Kit (Qiagen) according to manufacturer’s instructions. Genome-wide methylation analysis was performed on genomic DNA prepared from 12 colonic adenomas (3 mice for each of the four genotype-groups) and normal colonic mucosa from 6 C57BL/6 mice by standard procedures. Methylation levels were evaluated by Digital Restriction Enzyme Analysis of Methylation (DREAM), as previously reported (25). Briefly, methylation at specific CpG sites throughout the genome was evaluated by creating specific signatures at unmethylated and methylated CpG sites by sequential digests of genomic DNA with restriction endonucleases SmaI and XmaI, respectively. The restriction fragments with distinct signatures, 5′-GGG at unmethylated sites or 5′-CCGGG at methylated sites, were analyzed by next generation sequencing. Sequencing reads were mapped to the restriction sites in the reference genome, and signatures corresponding to methylation status of individual DNA molecules were resolved (26). Methylation ratios for each individual CCCGGG site were calculated as a proportion of methylated counts to the sum of unmethylated and methylated counts. Differences in methylation were evaluated by comparing the level of methylation in each genotype-group and the level of methylation in normal colonic mucosa from C57BL/6 mice.

### Analysis of DNA Methylation by Bisulfite Modification Sequencing

Bisulfite sequencing analysis was conducted to validate the difference in DNA methylation detected at the *Dirc2* and *Calb1* genes by DREAM and to investigate the DNA methylation of *Apc* promoter. Briefly, genomic DNA (250-500ng) was modified by sodium bisulfite reaction using the EZ DNA Methylation-Lightning kit (Zymo Research, Irvine, CA, USA) according to the manufacturer’s instructions. PCR amplicons were designed to verify the exact CpGs detected by the DREAM assay using Primer3 software (http://bioinfo.ut.ee/primer3-0.4.0/). A CpG cluster located in a 441bp region containing the *Apc* transcription site was analyzed according to Wang et al. (2016 Carcinogenesis). Products from the bisulfite reactions were amplified by PCR, purified using a PCR Purification Kit (QIAGEN) and sub cloned into pGEM T-Easy vector I (Promega). At least 12-19 individual clones were sequenced with the Sp6b reverse primer to determine the methylation status. Methylation levels were analyzed by QUMA software (http://quma.cdb.riken.jp/).

### RNA sequencing

A gene expression analysis by RNA-seq was conducted in triplicate per each genotype-group on the samples described previously in genome-wide methylation section. Briefly, RNA was isolated using TRIzol (Invitrogen, NY) according to the manufacturer’s protocol. The sequencing libraries were constructed from 500 ng of total RNA using the Illumina’s TruSeq RNA Sample pre kit V2 (Illumina) following the manufacturer’s instructions. The fragment size of RNAseq libraries was verified using the Agilent 2100 Bioanalyzer (Agilent) and the concentrations were determined using a Qubit instrument (LifeTech). The libraries were loaded onto an Illumina HiSeq 2500 at 6-10 pM on the rapid mode for 1×75bp single-end read sequencing. The fastq files were generated on the Illumina’s BaseSpace for further analysis.

### Quantitative real-time PCR

Validation of the RNA-seq data was performed in triplicate by quantitative real-time PCR (RT-PCR) for 8 selected loci (*Mmp9, Retnlg, Retnlb, Aldh1a3, Ccl4, Mptx1, Sprr2h* and *Chil3*) differentially expressed in *Tet1*^-/-^ *Tdg*^+/+^ *Apc*^Min/+^; *Tet1*^+/-^ *Tdg*^N151A/+^ *Apc*^Min/+^ and *Tet1*^+/+^ *Tdg*^N151A/+^ *Apc*^Min/+^ groups. RNA was reverse transcribed (RT) using Moloney murine leukemia virus reverse transcriptase (Ambion) and a mixture of anchored oligo-dT and random decamers (IDT). Taqman or SYBR Green assays were used in combination with Applied Biosystems Universal Master mixes and run on a 7900 HT sequence detection system (Applied Biosystems). Cycling conditions were 95°C, 15 min, followed by 40 (two-step) cycles (95°C, 15 s; 60°C, 60 s). Ct (cycle threshold) values were converted to quantities (in arbitrary units) using a standard curve (five points, four-fold dilutions) established with a calibrator sample. For each sample, the values are averaged and SD of data derived from at least two independent PCRs. Gene expression data was normalized to *Rpl32*. Primer sequences and conditions are available upon request.

## Results

### Adenoma multiplicity and characteristics in *Tet1* and/or *Tdg Apc*^*Min*^ mutant mice

To investigate the biological significance of the TET-TDG demethylation axis in intestinal tumorigenesis, we crossed *Tet1* knockout and/or mice bearing the dominant-negative *Tdg*^N151A^ allele with *Apc*^Min^ (Multiple Intestinal Neoplasia) mice, predisposed to intestinal adenomas (with a rare occurrence of cancers) and resembling the human familial adenomatous polyposis (FAP) condition (27). In particular, we used the *Apc*^Min^ Fox Chase Cancer Center variant characterized by high incidence of colonic adenomas (22). We studied a total of 90 mice with 4 genotypes: *Tet1*^*-/-*^*Apc*^Min/+^; *Tet1*^*+/-*^*Tdg*^*N151A/*+^*Apc*^Min/+^; *Tdg*^*N151A/*+^*Apc*^Min/+^ and *Tet1*^*+/+*^*Tdg*^*+/+*^ *(“wild type”) Apc*^Min/+^. We first investigated whether *Tet1* and /or *Tdg* defects modify tumor load (tumor multiplicity and size) in *Apc*^Min^ background. Through a detailed inspection of the gastrointestinal tract, we found that *Tet1*^*-/-*^ knockout and *Tet1*^*+/-*^*Tdg*^*N151A/*+^ double heterozygotes exhibited an increased number of large colonic adenomas (> 3-7mm) compared with *“wild type” Apc*^Min/+^ mice (Fig. 1A-D).

**Fig 1.**
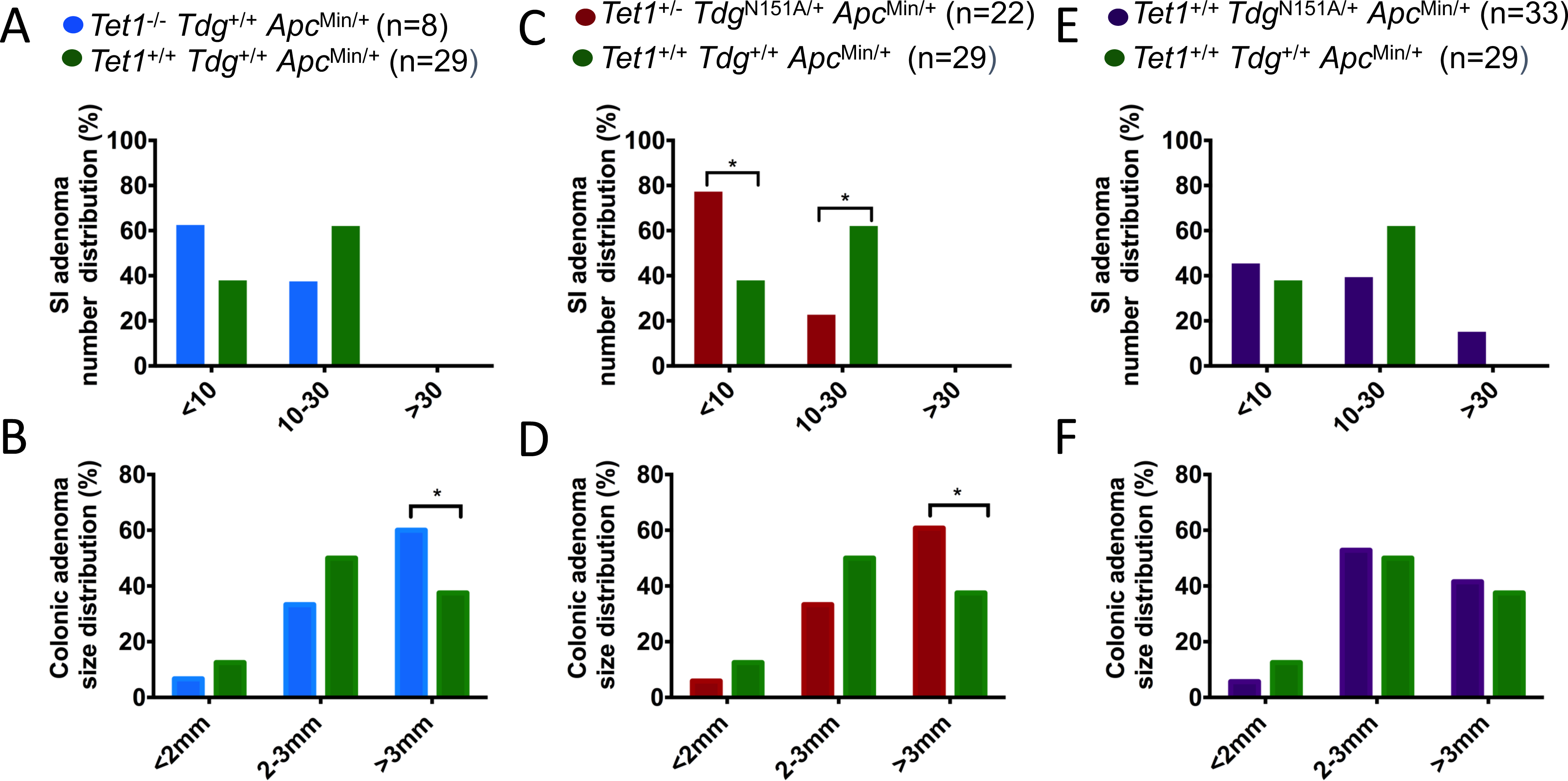
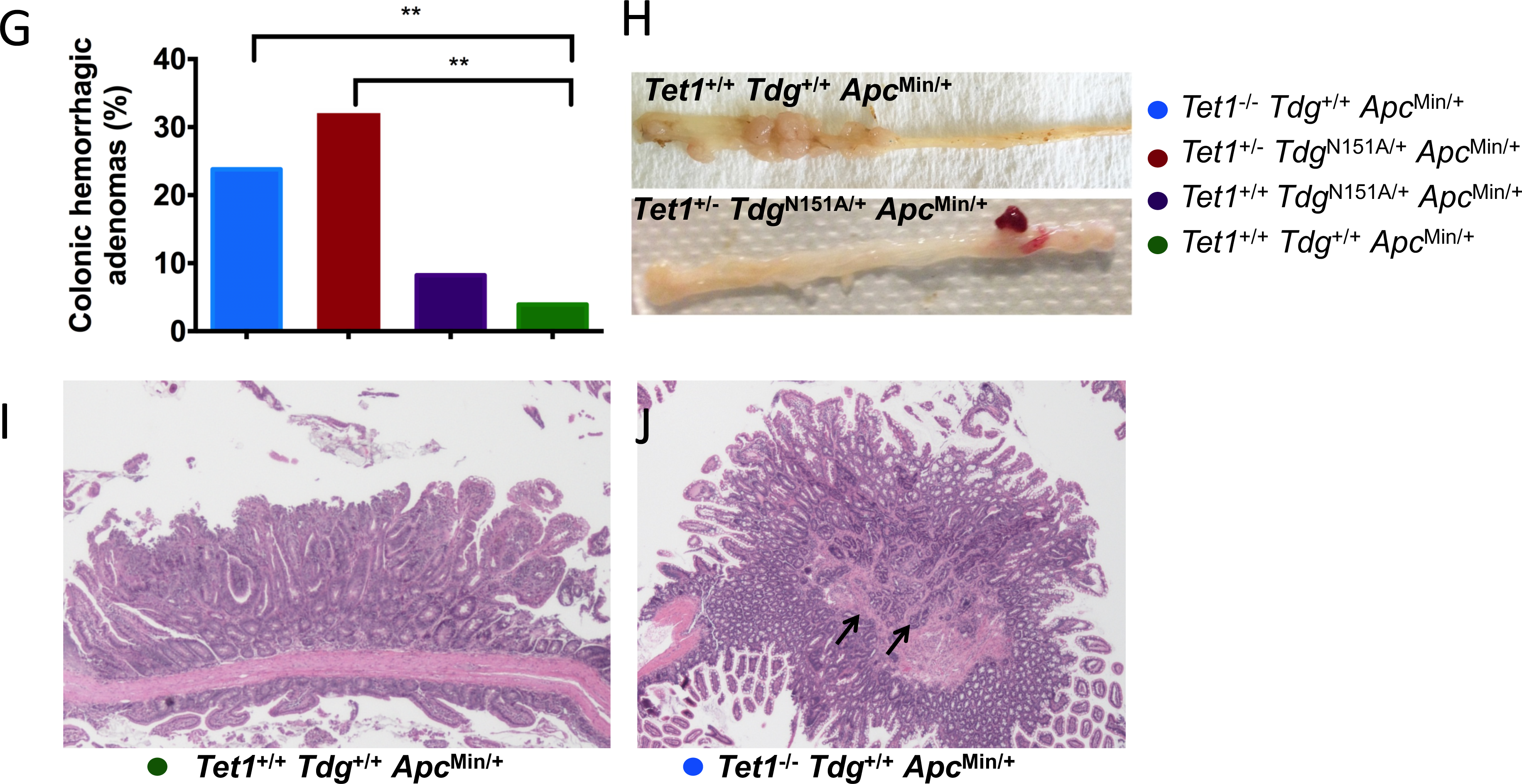
*Tet1* and *Tdg* inactivation enhances tumorigenesis in *Apc*^**Min/+**^ background. Graphical representation of the total number and size distribution of **(A, C, E)** small intestinal, and **(B, D, F)** colonic adenomas in *Tet1*^*-/-*^, *Tet1*^*+/-*^ *Tdg*^*N151A/+*^, *Tdg*^*N151A/+*^ and “wild type” *Apc*^Min/+^ mice. **(G)** Percent of hemorrhagic colonic adenomas in the four genotype-groups. * *p*-value =0.0044, ** *p*-value =0.0008. **(H)** Representative pictures of colons of *Apc*^Min/+^ (top panel) and *Tet1*^*+/-*^ *Tdg*^*N151A/+*^ mice (bottom panel). **(I, J)** Hematoxylin & Eosin-stained sections of colonic adenomas developed in *Tet1*^*-/-*^, and “wild type” *Apc*^Min/+^.

We also found that a fraction of *Tdg*^N151A/+^ single heterozygous mice showed a predisposition to develop an increased number of intestinal adenomas (>30), mostly in the small intestine. This phenotype was not observed in *Tet1*^*-/-*^ homozygotes, *Tet1*^*+/-*^*Tdg*^*N151A/*+^ double heterozygotes or *“wild type”Apc*^Min/+^ mice (Fig. 1E-F).

Remarkably, a statistically significant increase of congested, hemorrhagic adenomas was observed in *Tet1*^*-/-*^ homozygotes (*p*-value =0.0044) and *Tet1*^*+/-*^ *Tdg*^*N151A/*+^ double heterozygotes (*p*-value =0.0008) compared with *“wild type” Apc*^Min/+^ mice (Fig. 1G-H). Of note, the hemorrhagic adenomas in *Tet1*^*-/-*^ homozygotes and *Tet1*^*+/-*^ *Tdg*^*N151A/*+^ double heterozygotes mice were limited to the colon and were rarely detected in *“wild type”Apc*^Min/+^ mice. Histopathological analysis of hemorrhagic adenomas from *Tet1*^*-/-*^ homozygotes and *Tet1*^*+/-*^*Tdg*^*N151A/*+^ double heterozygotes revealed features of invasion compared with *“wild type”Apc*^Min/+^ mice (Fig. 1I-J). Incidentally, no other tumors outside the digestive tract and no significant differences in survival were observed in *Tet1* and/or *Tdg* deficient mice compared with *“wild type”Apc*^Min/+^ mice.

These observations indicate that both *Tdg* and *Tet1* act as tumor suppressor genes in the intestine, affecting tumor size (*Tet1*) or multiplicity (*Tdg*). In addition, homozygous or heterozygous *Tet1* mutations enhance intestinal tumorigenesis in the *Apc*^Min/+^ background by causing hemorrhagic adenomas with features of invasion (incipient conversion to malignancy).

### Genome-wide DNA methylation analysis

The higher tumor load observed in *Tet1* and *Tdg* mutant compared with *“wild type”Apc*^Min/+^ mice suggests that *Tet1* and *Tdg* inactivation may have an impact on the intestinal tumorigenesis by an aberrant modulation of the epigenome, due their crucial role in active DNA demethylation. Our goal was to identify the epigenomic changes associated with *Tet1* and *Tdg* defects in the context of *Apc*^Min/+^ mutation. To this purpose, we conducted a genome-wide DNA methylation analysis by digital restriction enzyme analysis of methylation (DREAM)(26), which allows for identification of differentially methylated sites in CpG islands (CGI) and non-CpG islands (NCGI) at high resolution (it interrogates approximately 25K CpG sites in the murine genome). DREAM analysis was conducted on triplicate adenomas from mice within the four genotype groups, and on triplicate samples of colonic mucosa from C57BL6 mice.

We also investigated the differences in DNA methylation between *Tet1*^*-/-*^, *Tdg*^*N151A/*+^, *Tet1*^*+/-*^*Tdg*^*N151A/*+^ and “wild type” *Apc*^Min/+^ adenomas more systematically by analyzing CGI and NCGI sites separately. Compared to normal colonic mucosa, a global hypomethylation at NCGIs was detected in *Apc*^Min/+^ adenomas and corresponds to the genome-wide hypomethylation reported in human colonic adenomas as one of the earliest manifestations of multistep tumorigenesis in the colon (28-30)(Fig. 2A). Remarkably, this global hypomethylation was progressively reduced in *Tdg*^*N151A/*+^, *Tet1*^*+/-*^ *Tdg*^*N151A/*+^ and *Tet1*^*-/-*^ adenomas (Fig. 2A), which indicates a shift towards increased methylation levels in the absence of a functional TET-TDG axis. Previous DNA methylation analysis of Apc^Min^ tumors showed that these do not have much CpG island hypermethylation or CIMP (31). Interestingly, we found that only *Tet1*^*-/-*^ adenomas showed elevated methylation at many CGI sites (Fig. 2B), which resembles the CpG island methylator phenotype (CIMP) in humans. No microsatellite instability (MSI), a phenotype associated with CIMP in human colorectal cancers, was detected in *Tet1* deficient colonic adenomas (data not shown). Overall, a shift towards increased methylation was detected when Tet1 and Tdg mutant adenomas were compared to “wild type” Apc-Min adenomas (Fig. 2C).

**Fig 2.**
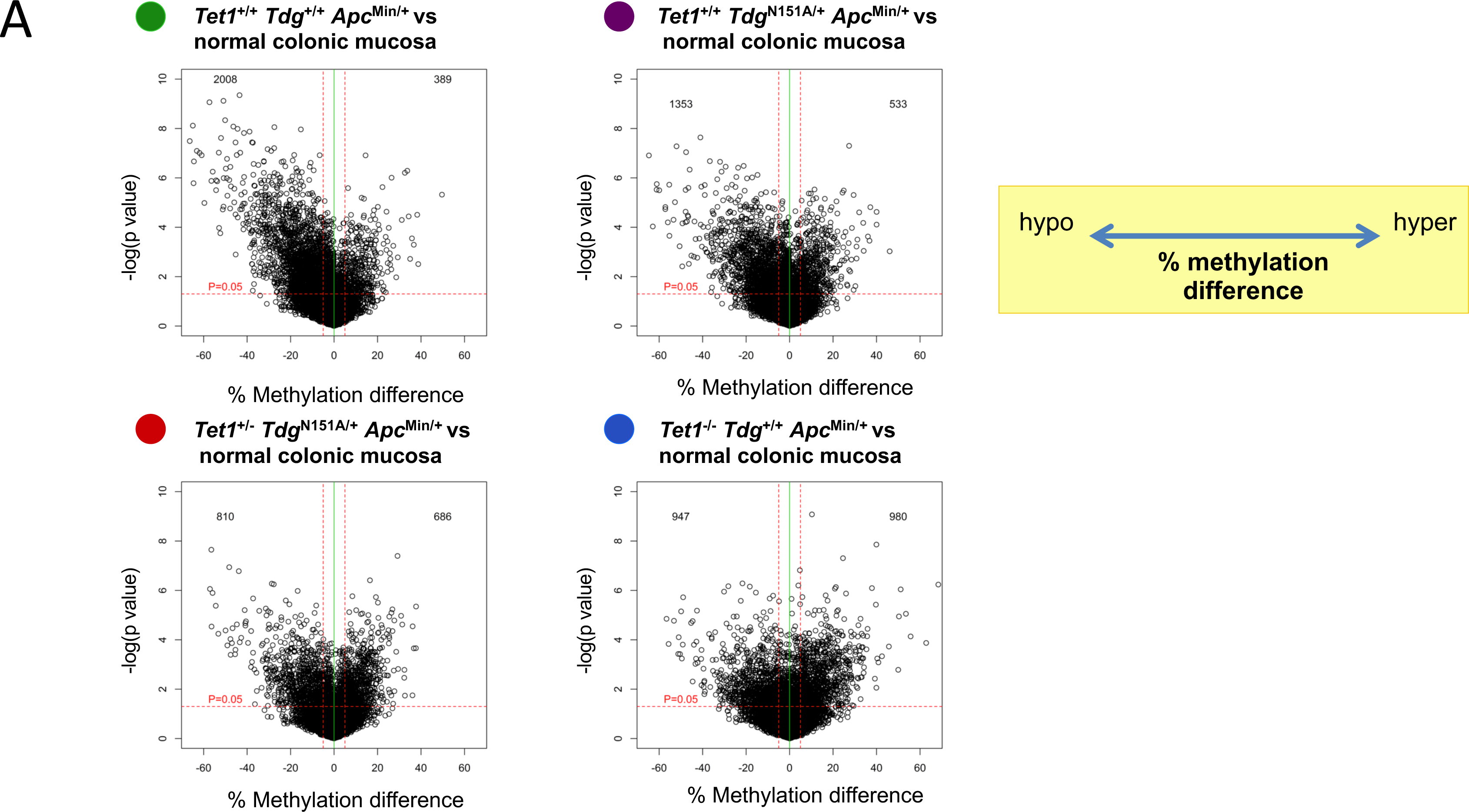
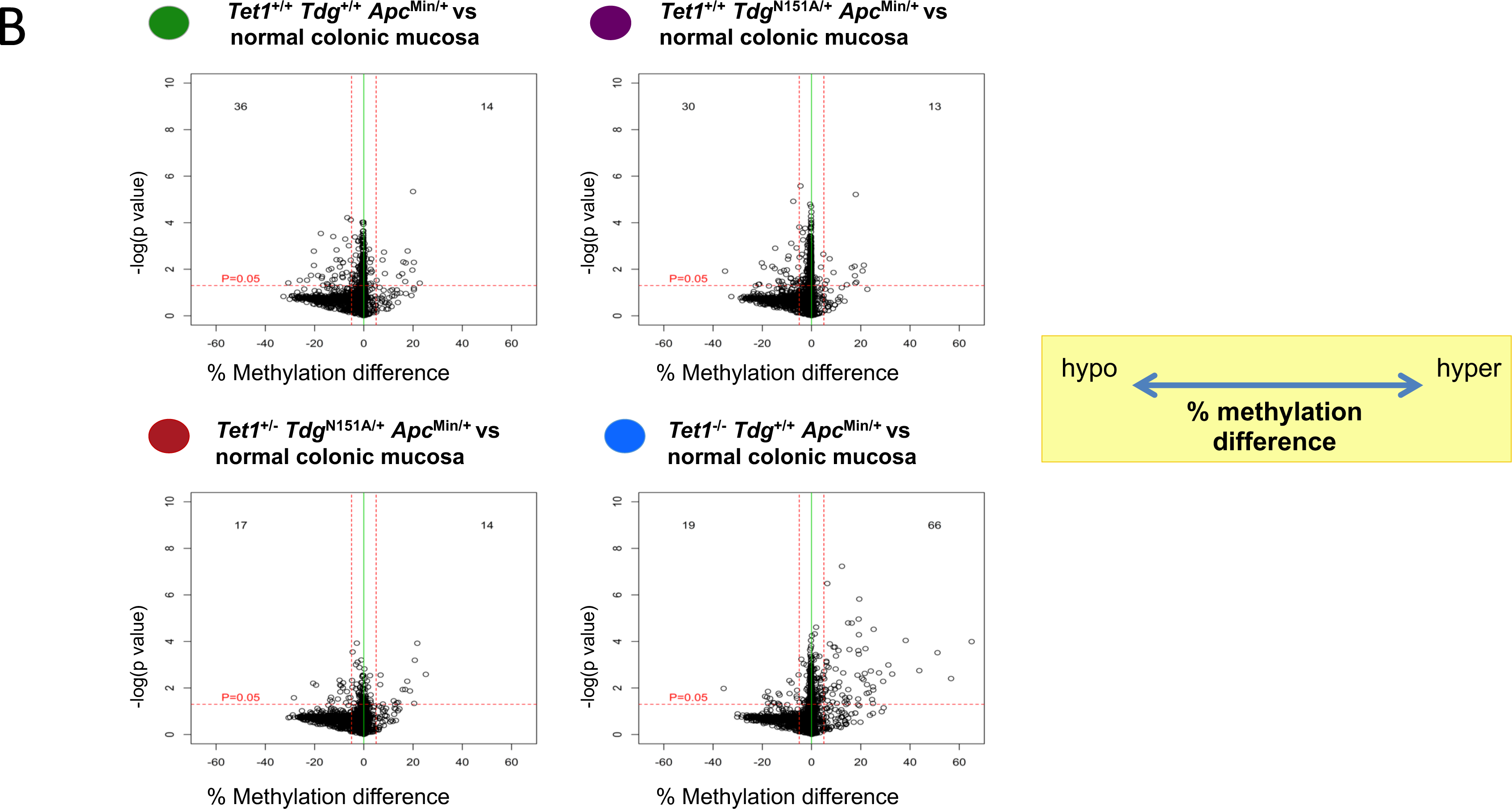
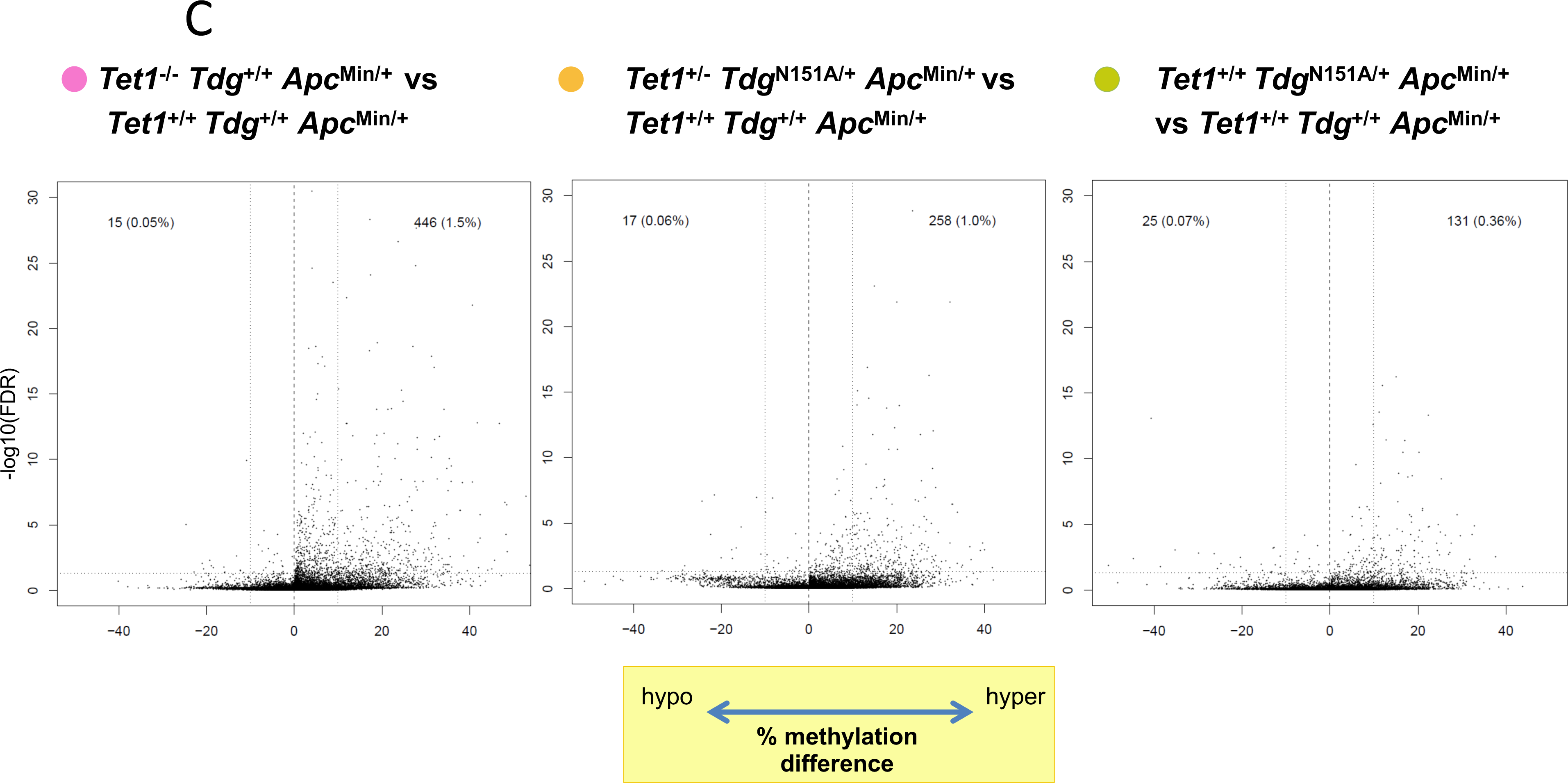
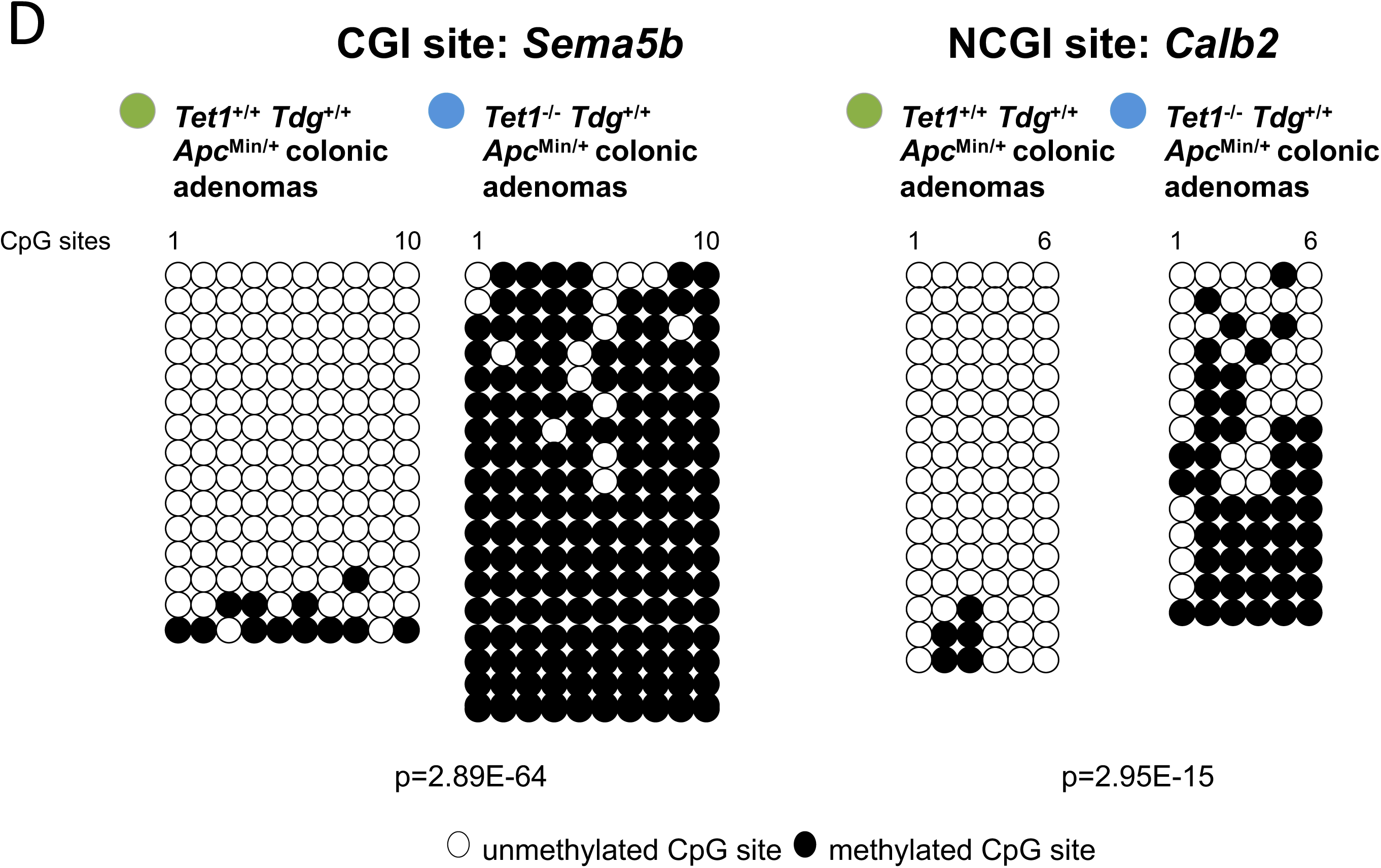
Methylome analysis shows loss of global hypomethylation in *Tet1*- and *Tdg*-mutant adenomas. Volcano plots showing the difference in DNA methylation at (A) non-CpG island (NCGI) and (B) CpG island sites (CGI) in each comparison. The average of methylation of *Tet1*- and/or *Tdg*-mutant and *Apc*^Min/+^ control adenomas versus normal colonic mucosa and the p value for each site are reported on the x-axis and y-axis respectively. **(C)** Volcano plots showing the global difference in DNA methylation in each comparison between adenomas of different genotype. **(D)** Example of DNA methylation levels validation for a CGI (*Semaf5b*) and a NCGI (*Calb2*) sites differentially methylated in *Apc*^Min/+^ and *Tet1*^-/-^ colonic adenomas by bisulfite sequencing.

Next, we confirmed the methylation differences by bisulfite sequencing for two selected genes, *Sema5b* and *Calb2*, which were hypermethylated in *Tet1*^*-/-*^ colonic adenomas at CGI and NCGI sites, respectively. We investigated the genomic DNA extracted from 3 *Tet1*^*-/-*^ and 3 *Apc*^Min/+^ adenomas belonging to 6 different mice. 47 and 12 CpG sites were analyzed for *Sema5b* and *Calb2*, respectively. Analysis of these CpG sites confirmed hypermethylation in all the 3 *Tet1*^*-/-*^ adenomas. Fig. 2D shows an overview of the significant increase in methylation at CpG sites for *Sema5b* (*p*- value =2.89E-64) and *Calb2* (*p*-value =2.95E-15) in the 3 *Tet1*^*-/-*^ samples analyzed compared to “wild type” *Apc*^Min/+^ samples.

### Genome-wide transcriptomic analysis

To clarify the role of *Tet1* and *Tdg* in colorectal tumorigenesis and evaluate the impact of the methylation changes observed in *Tet1* and *Tdg-*deficient adenomas on gene expression, we conducted a genome-wide transcriptomic analysis on *Tet1*^*-/-*^, *Tet1*^*+/-*^*Tdg*^*N151A/*+^, *Tdg*^*N151A*^ and *Apc*^Min/+^ adenomas. Using a FDR cutoff of 5%, a total of 224 genes were found differentially expressed between the three mutant genotype-groups and the *Apc*^Min/+^ control group. Gene ontology analysis conducted on the 224 differentially expressed genes revealed an enrichment of inflammation, innate immunity, interferon signaling and antimicrobial response processes (Fig. 3). A comparative analysis between each mutant genotype-group and *Apc*^Min/+^ control group identified 88, 74 and 123 genes differentially expressed in *Tet1*^*-/-*^, *Tet1*^*+/-*^*Tdg*^*N151A/*+^, *Tdg*^*N151A/*+^ adenomas, respectively. We observed that 10 genes were shared among the three mutant genotype-groups, including calcium- and zinc-binding proteins (*S100A8* and *S100A9*) implicated in differentiation of stromal myeloid cells (neutrophils and macrophages) and cytokine production (*Exp. Med. 205 (10): 2235*–*49)*, hormones (*Retnlb* and *Retnlg*) and an antimicrobial peptide (*Defb45*) involved in myeloid dendritic cell chemotaxis and antimicrobial response, respectively (Fig. 3a). We also compared the differentially expressed genes between *Tet1*^*-/-*^ and *Tdg*^*N151A/*+^, *Tet1*^*+/-*^*Tdg*^*N151A/*+^ and *Tdg*^*N151A/*+^ and *Tet1*^*-/-*^ and *Tet1*^*+/-*^*Tdg*^*N151A/+*^ genotype-groups. GO analysis of the genes shared only between *Tet1*^*-/-*^ and *Tdg*^*N151A/*+^ (n=10), *Tdg*^*N151A/*+^ and *Tet1*^*+/-*^*Tdg*^*N151A/*+^ (n=26) revealed that they were mostly immunoglobulins (*Ighg1, Ighg2b, Igkv6-15* and *Igkv1-117*) and chemokines or their receptors (*Cxlc2, Ccl3, Nos2, Cxcl3, Osm, Csf3, Il1r2, Chi3l3* and *Arg1*), respectively. No enrichment of biological processes was detected by GO analysis of the five genes (*Snhg11, Mptx1, Aldh1a3, AI747448* and *Adh1*) in common between *Tet1*^*-/-*^ and *Tet1*^*+/-*^*Tdg*^*N151A/+*^ genotype-groups.

**Fig 3.**
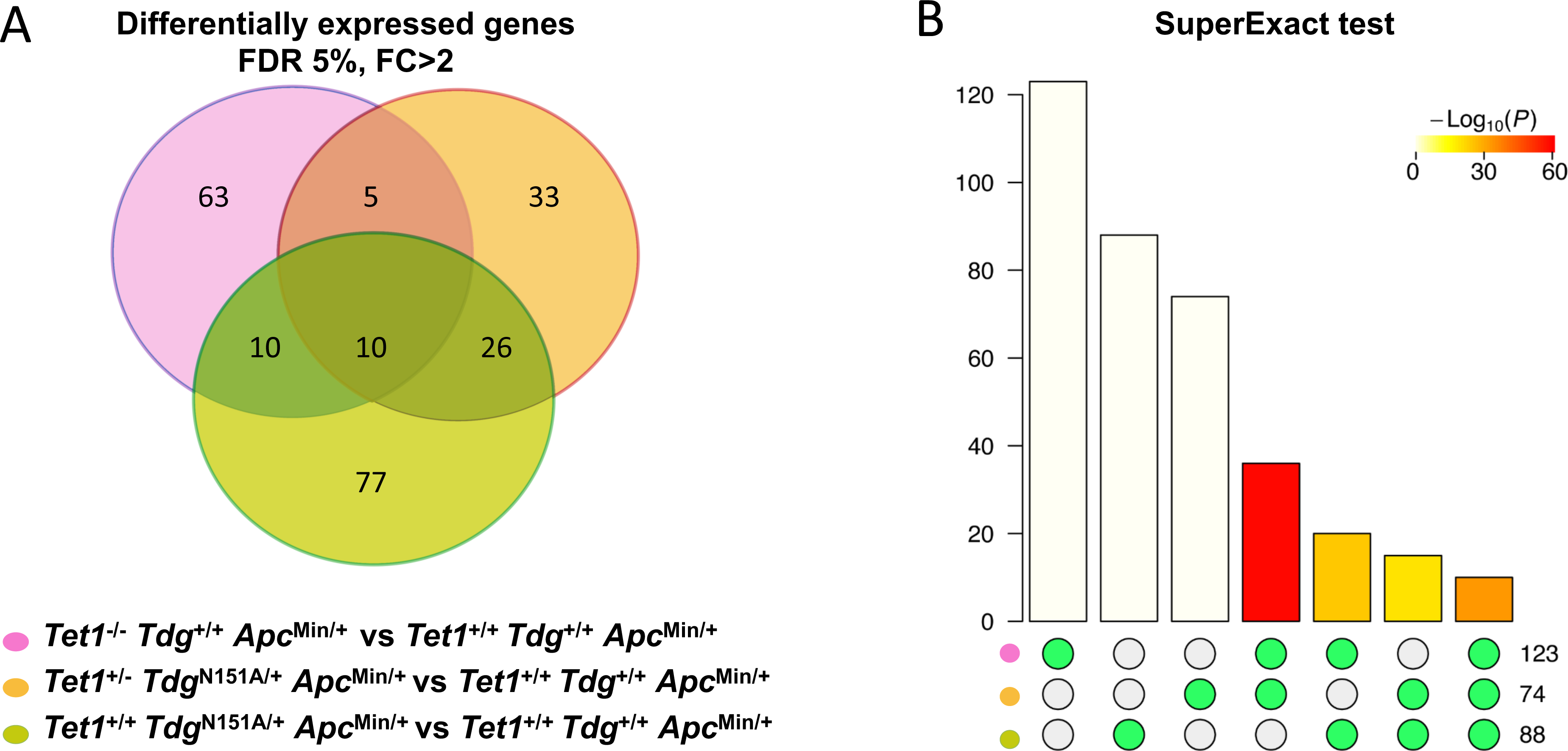
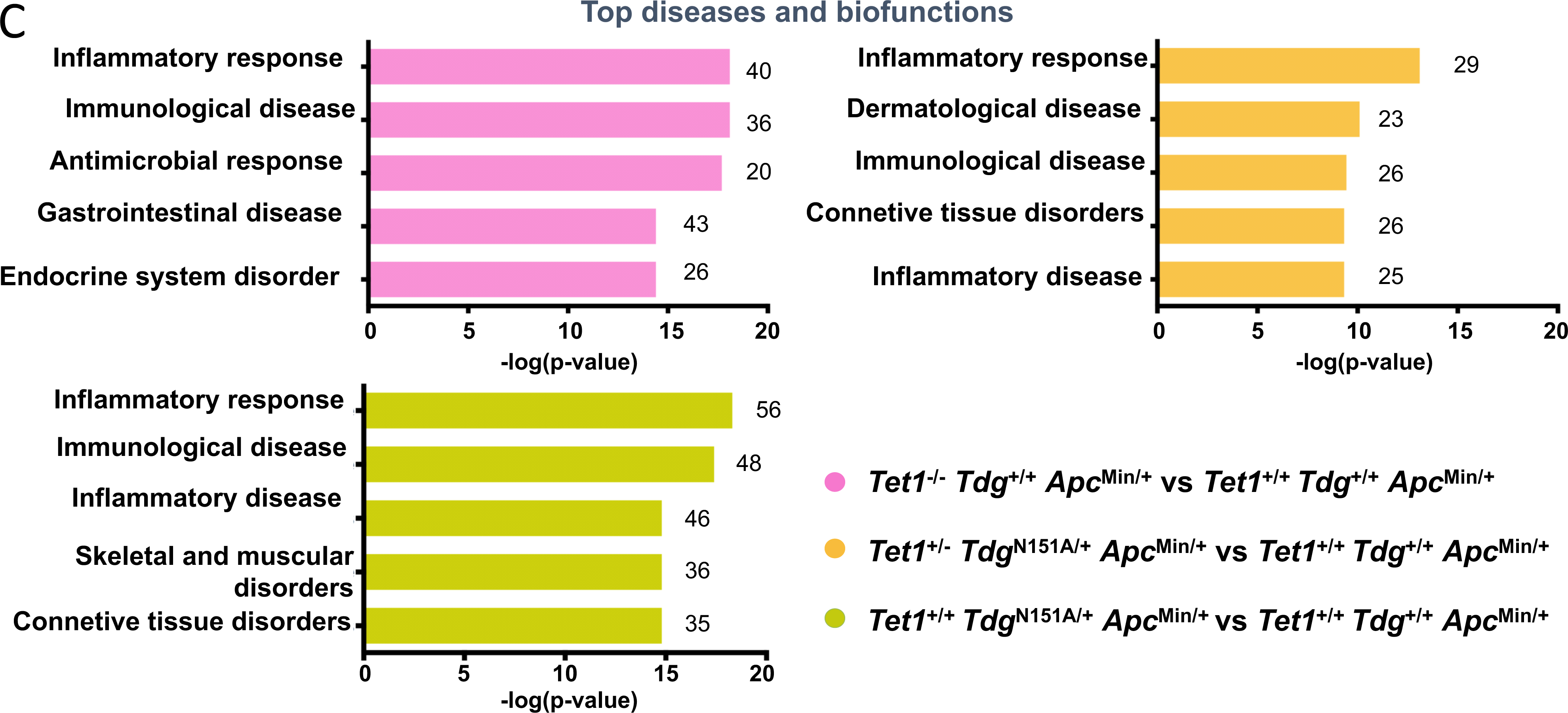
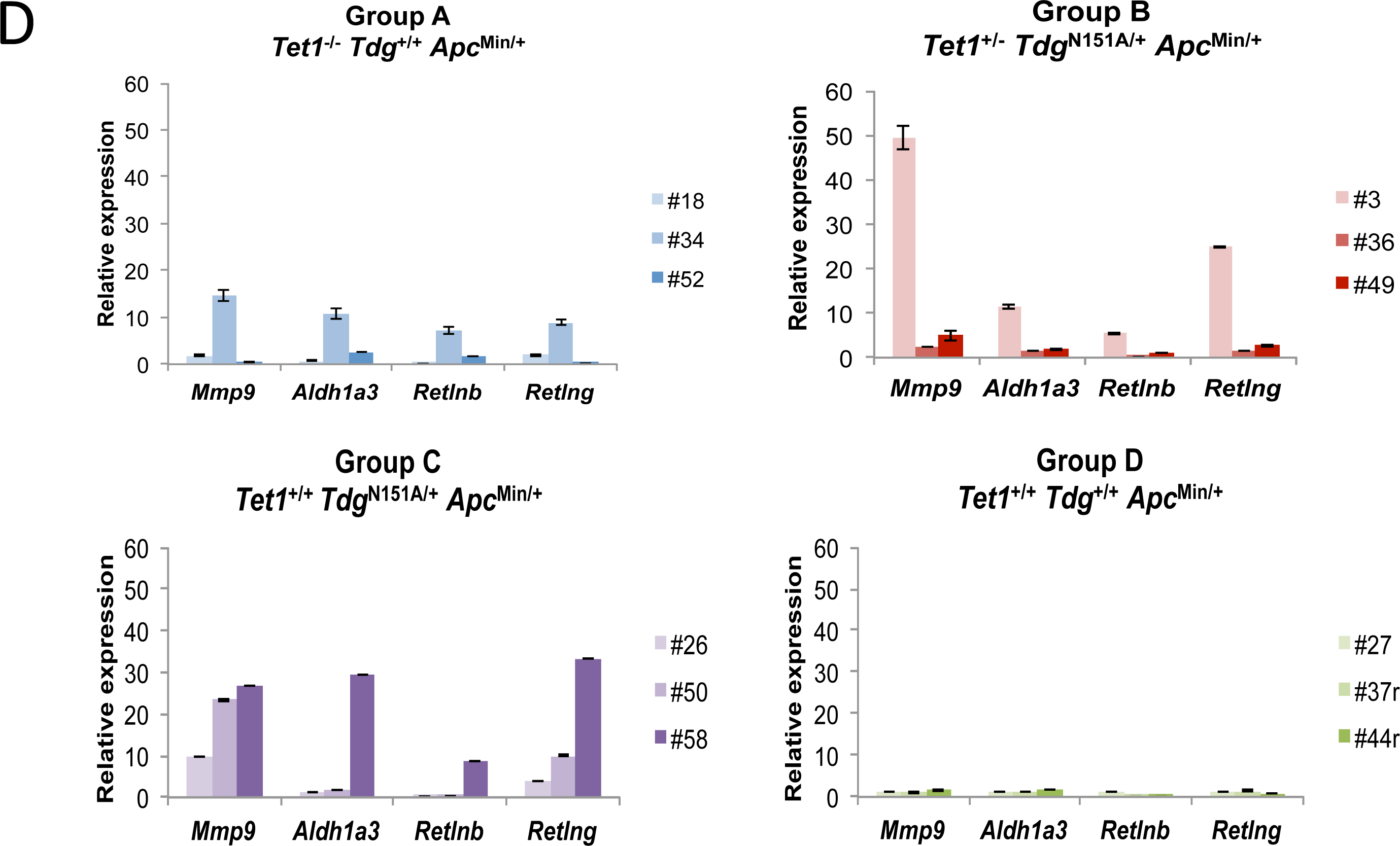
*Tet1*- and/or *Tdg*-mutant adenomas exhibit upregulation of inflammatory and immune response pathways. **(A)** Venn diagrams showing the number of common and differentially expressed genes (FDR 5% and FC>2 relative to *Apc*^Min/+^ controls) in *Tet1*^*-/-*^, *Tdg*^*N151A/+*^ and *Tet1*^*+/-*^ *Tdg*^*N151A/+*^ adenomas. (**B**) A bar chart illustrating all possible intersections by Super-exact test among genes differentially expressed in *Tet1*^*-/-*^, *Tdg*^*N151A/+*^ and *Tet1*^*+/-*^ *Tdg*^*N151A/+*^ adenomas confirms statistically significant involvment of the same genes in Tet1 and Tdg mutant adenomas. The matrix of solid and empty circles at the bottom illustrates the “presence” (solid green) or “absence” (empty) of the data sets in each intersection. The numbers to the right of the matrix are set sizes. The colored bars represent the intersection sizes with the color intensity showing the P value significance. **(C)** Ingenuity pathway analysis of the upregulated genes in *Tet1*^-/-^, *Tdg*^*N151A/+*^ and *Tet1*^*+/-*^ *Tdg*^*N151A/+*^ adenomas. The most highly enriched gene ontology categories (top diseases and bio functions) are shown. **(D)** Example of gene expression validation for two selected differentially expressed genes in *Tet1*^*-/-*^, *Tdg*^*N151A/+*^ and *Tet1*^*+/-*^ *Tdg*^*N151A/+*^ adenomas by RT–qPCR. Data were normalized to *Rpl32* control gene. Standard deviations were shown with the error bars.

We also identified genes specifically expressed in each mutant genotype-group. In particular, 77 genes differentially expressed only in *Tdg*^*N151A/*+^ adenomas were mainly interleukins (*Il1b, Il1a, Il1f9* and *Il18rap)*, chemokines (*Chi3l1, Ccrl2, Ccl4, Ccr1, Clca2, Cxcr2, Clec4e, Clec4d* and *Irg1*) and immunoglobulins (*Ighg2c, Ighv5-6, Ighv1-19, Ighv1-81, Igkv10-96, Igkv14-111* and *Igkv1-135)*. Few genes belonging to the TRIM family proteins (*Trim10, Trim15, Trim29)* induced by the interferons were found also specifically expressed in *Tdg*^*N151A/*+^ adenomas. Conversely, most of the 63 genes differentially expressed in *Tet1*^*-/-*^ knockouts mice are involved in interferon signaling (*Oasl1, Oasl2, Oas3, Oas1a, Itln1, Itih5, Isg15, Ifit1, Ifit3, Ifit44, Trim30a, Rtp4* and *Rsad2*), major histocompatility complex (*H2-Q5, H2-Q6, H2-Q7, H2-Aa, H2-Ab1, H2-Eb1)* and the antimicrobial defense response (*Defa3, Defa21, Defa22, Defa24* and *Liz1*). No cytokines and few immunoglobulins *(Igkc, Iglv1* and *Iglc2)* were found specifically expressed in *Tet1*^*-/-*^ knockouts mice. None of the 33 genes with specific expression in *Tet1*^*+/-*^*Tdg*^*N151A/*+^ adenomas showed an enrichment of cytokines, interferon signaling or antimicrobial defense genes or other biological processes, except the presence of few immunoglobulins (*Ighv9-1, Igkv3-2, Igkv4-53, Igkv4-68* and *Igkv5-39)*.

Importantly, GO analysis also revealed a distinct gene expression profiles in *Tet1*^*-/-*^ knockouts and *Tdg*^*N151A/*+^ single heterozygotes mice. In particular, *Tet1*^*-/-*^ knockouts mice showed a deregulation of interferon signaling and antimicrobial defense response, *Tdg*^*N151A/*+^ single heterozygotes mice an upregulation of inflammatory response. Moreover, we also observed that three mutant genotype-groups exhibited a similar over-expression profile, characterized by the upregulation of genes implicated in immune response.

Next, using pathway analysis we confirmed that the gene expression changes associated with *Tet1* and /or *Tdg* defects were involved in inflammatory response and immunological disease. We also found diseases/bio functions specifically associated with each mutant genotype-group. In particular, antimicrobial response, endocrine system disorders and gastrointestinal diseases were found in *Tet1*^*-/-*^ knockouts, dermatological disease and conditions in *Tet1*^*+/-*^*Tdg*^*N151A/*+^ double heterozygotes and skeletal and muscular disorders in *Tdg*^*N151A/*+^ single heterozygotes. Moreover, connective tissue disorders and inflammatory disease were in common among *Tet1*^*+/-*^*Tdg*^*N151A/*+^ double and *Tdg*^*N151A/*+^ single heterozygotes (Fig. 3)

Finally, we validated eight selected differentially expressed genes (*Mmp9, Retnlg, Retnlb, Aldh1a3, Ccl4, Mptx1, Sprr2h* and *Chil3)* in the three mutant genotype groups using quantitative RT-PCR (Fig. 3C and data not shown).

### Histopathology

We conducted histopathological analysis to identify morphological correlates of the activation of immune response and inflammatory pathways. This analysis revealed increased surface erosion, stroma activation (desmoplasia) and inflammation in adenomas from *Tet1*^*-/-*^, *Tet1*^*+/-*^*Tdg*^*N151A/*+^ double heterozygous and *Tdg*^*N151A/*+^ mice, in comparison with “wild type” *Apc*^Min/+^ mice (Fig. 4A). In addition, immunohistochemical staining with the f4/80 antibody revealed that Tet1- and/or Tdg-mutant adenomas show an increased infiltration of macrophages in the stroma surrounding the adenomatous crypts, but not morphologically normal crypts (Fig. 4B). No differences in T lymphocyte infiltration (as detected by CD3 staining) were found among adenomas of the four genotypes (data not shown).

**Figure 4.**
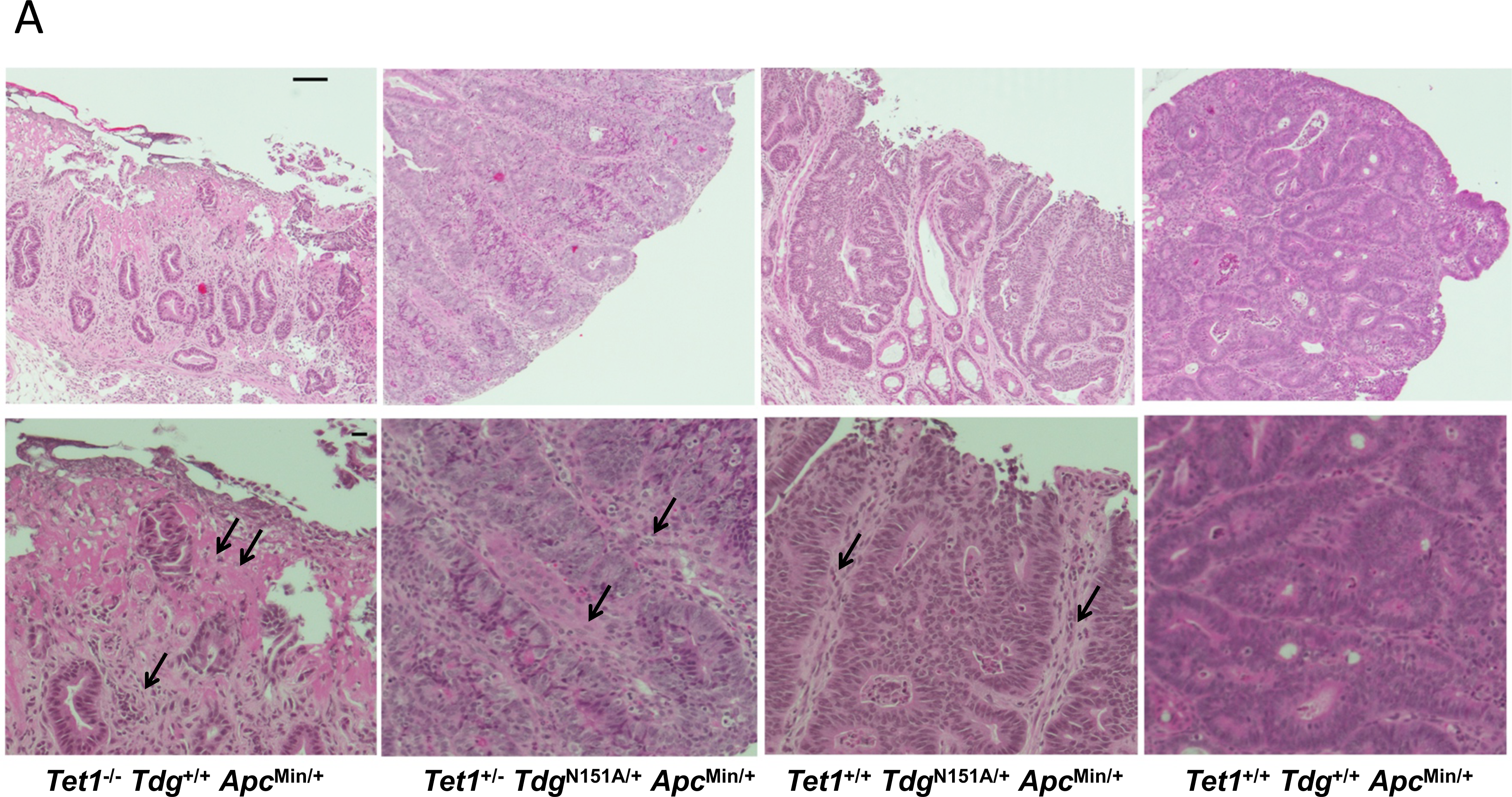
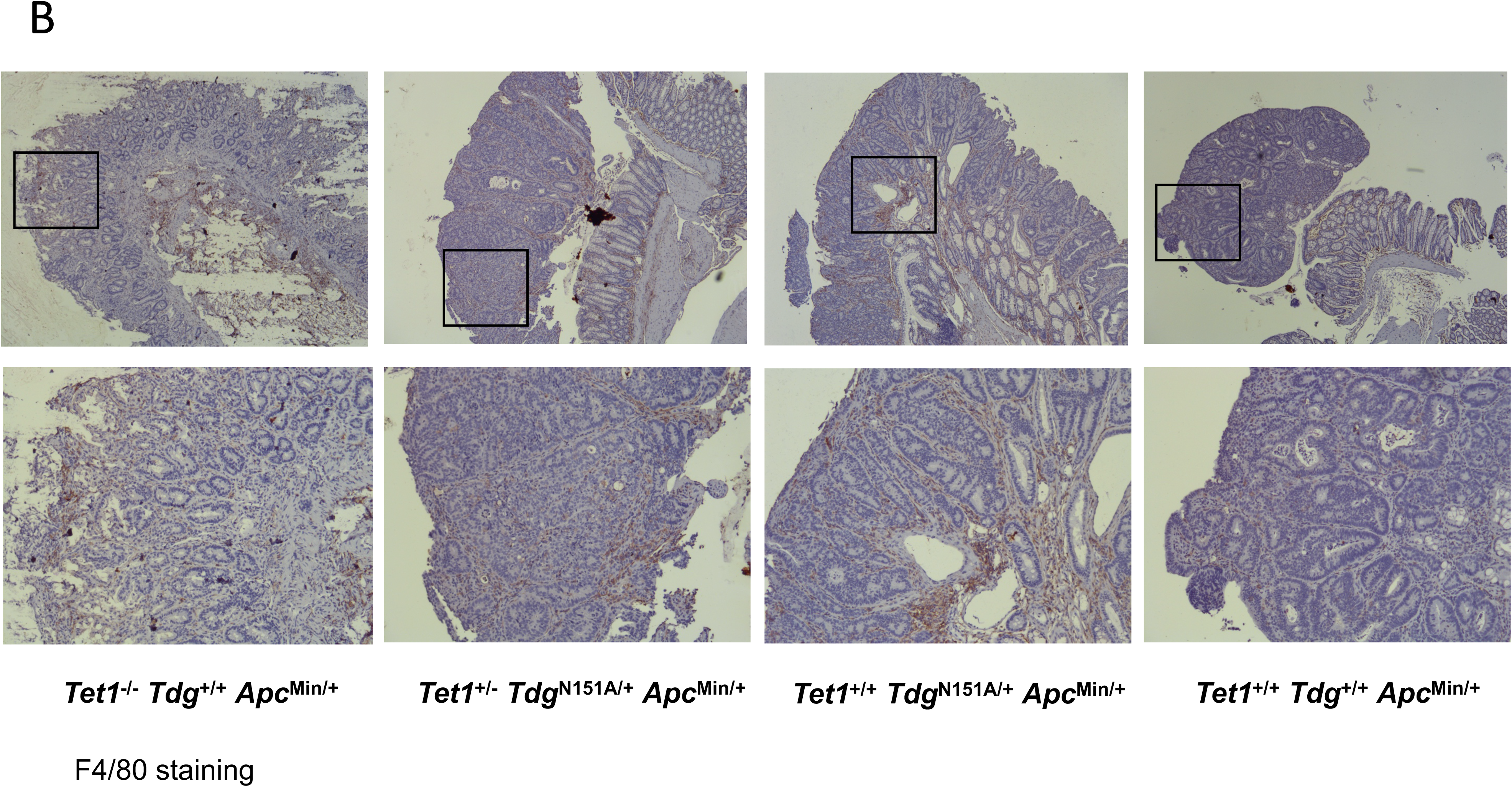
Histopathological analysis of adenomas of different genotypes stained by hematoxylin-eoson (**B**) or by immunohistochemistry with anti-f4/80 antibody (**B**).

### Identification of a Tet1- and Tdg-related inflammatory signature; significance in human CRC

We conducted an unsupervised hierarchical clustering analysis for RNA expression levels for differentially expressed genes. Three samples (A34, B3 and C58) with *Tet1* and/or *Tdg* defects clustered together and identified a list of 169 highly divergent, inflammatory genes (Fig. 5A). Remarkably, A34, B3 and C58 samples shared the upregulation of 75 genes mostly implicated in inflammation and innate immunity. Specifically, most of the 75 genes were interleukins (*Il-11* and *Il-1a*), chemokines (*Cxcl2, Cxcl3, Ccl4* and *Ccl3*), cytokine receptors (*Il1-r2* and *Cxcr2)*, lectins (*Clec4e, Clec4d)*, receptors (*Csf3r, Fpr1* and *Fpr2)*, calcium- and zinc-binding protein (*S100A8* and *S100A9)*, hormone (*Retnlg)* expressed mainly by macrophages and other myeloid cells. Importantly, the corresponding human 135-gene signature separates the colon adenocarcinoma (COAD) TCGA samples in four clusters, that align strongly with their microsatellite stable (MSS/CIN) or unstable status (MSI/MIN)(p<0.0001)(Fig. 5B). Whereas cluster 1 has mixed MSI/MSS status and cluster 3 corresponds to MSI cases, clusters 2 and 4 which are classified as MSS/CIN, display low and high inflammation gene expression signature, respectively (Fig. 5B). Importantly, cluster 4 (high inflammatory signature, *“hot”*) has low TET1 expression and vice versa cluster 2 (low inflammatory signature, *“cold”*) has high TET1 expression. TDG expression did not vary across the four clusters, with the caveat that we are not measuring its activity. This suggests that the (epi)genetic program set forth by TET1-TDG suppresses inflammation; and its alterations in human CRC are strongly linked to, and help stratify, the intrinsic features of CIN.

**Figure 5.**
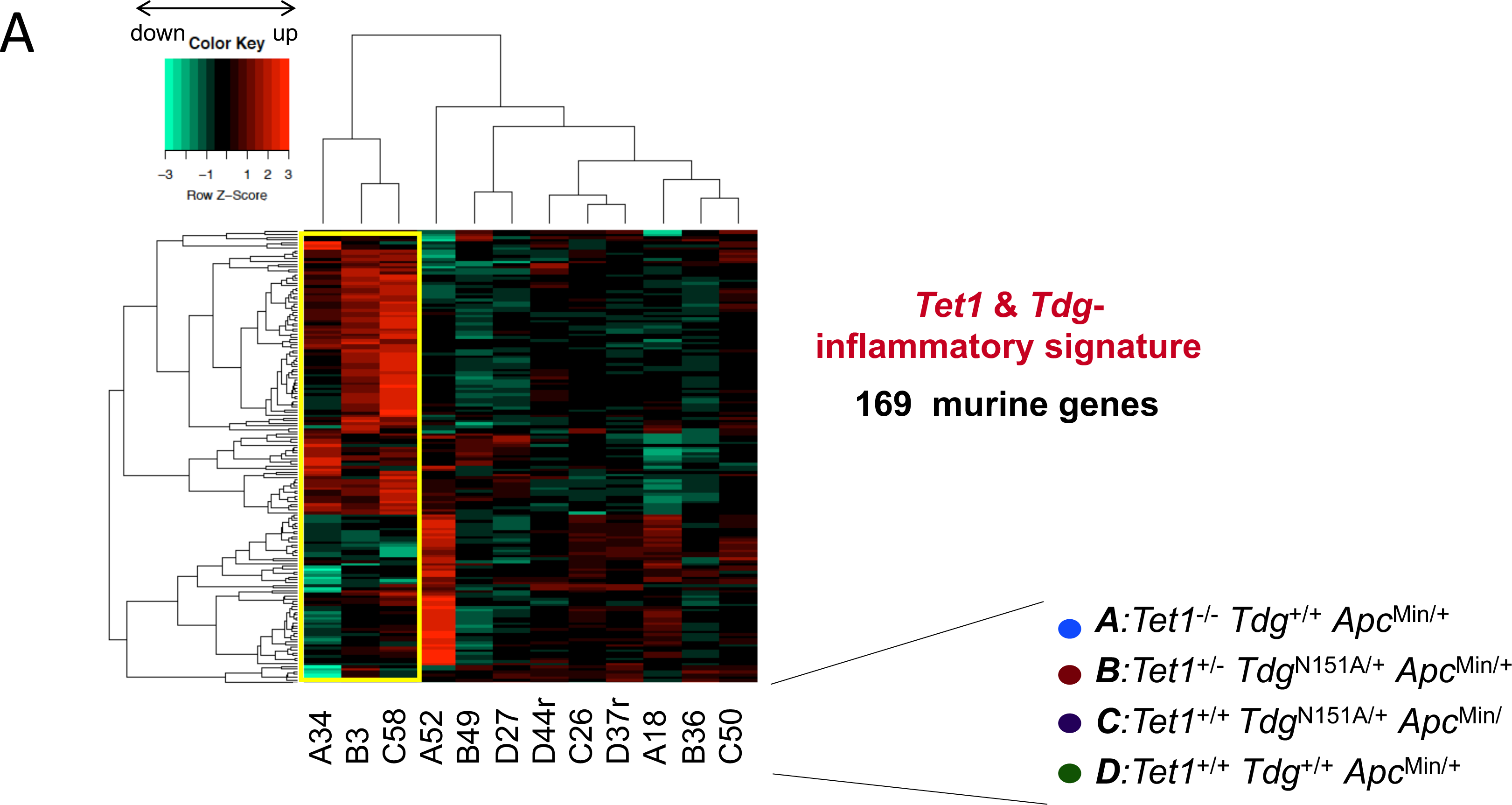
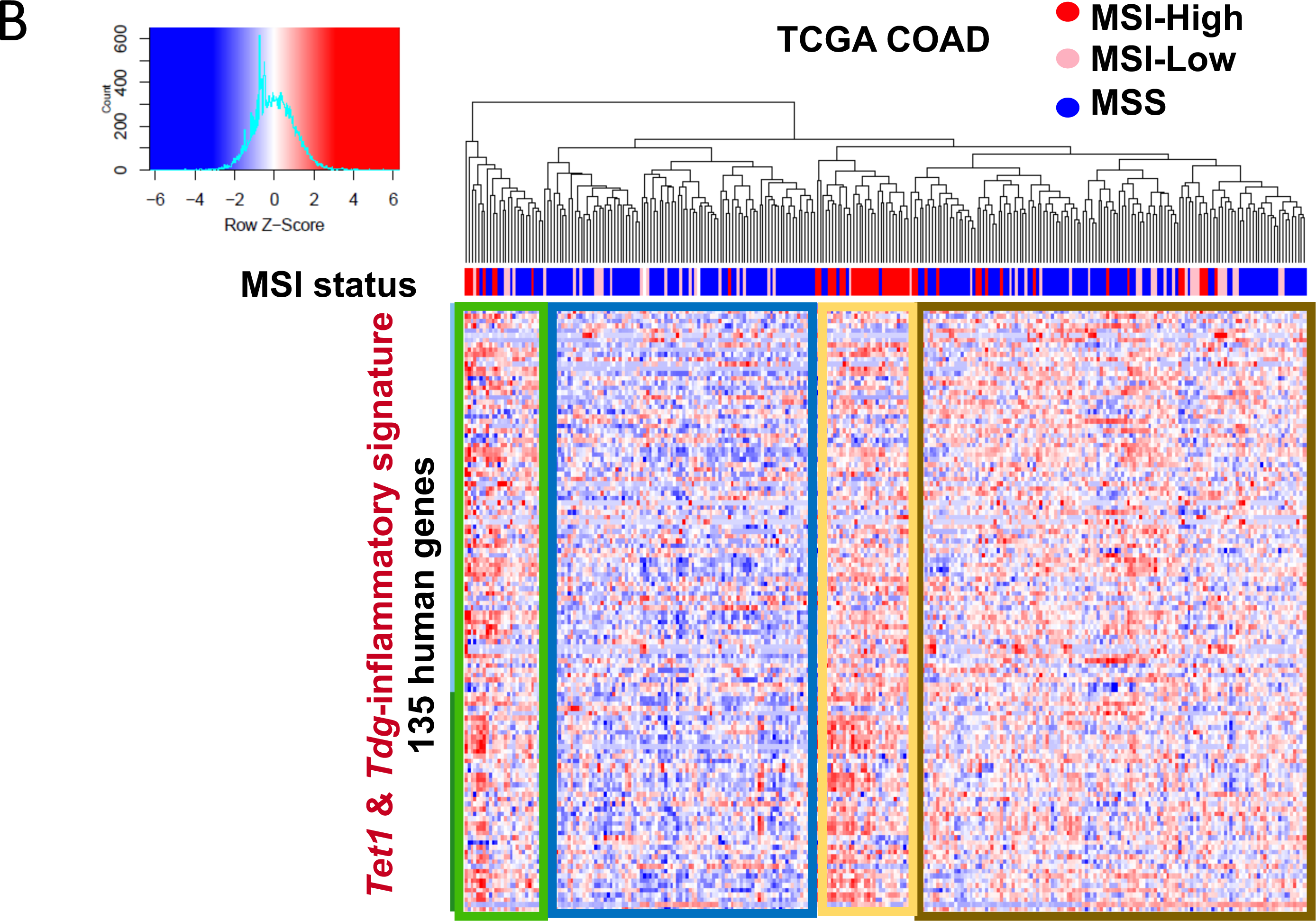
**(A)** Unsupervised hierarchical clustering analyses for RNA expression levels of 12 adenoma samples from the four genotypes (triplicates for each genotype) leads to the identification of a 169-gene inflammatory signature. (**B**) Tet1 & Tdg-inflammatory signature identifies 4 groups in TCGA COAD (colon adenocarcinoma) based on microsatellite status: cluster analysis of TCGA COAD samples with the corresponding human 135-gene inflammatory signature.

## Discussion

In this article, we show that both TET1 and TDG act as tumor suppressors in colorectal tumorigenesis, by affecting the number (TDG) or the size (TET1) of colonic adenomas. In addition, TET1 is involved in progression to malignancy, because colonic adenomas from *Tet1*^-/-^ and *Tet1*^+/-^*Tdg*^N151A/+^ mice show congestive/hemorrhagic appearance, corresponding to morphological features of invasion and incipient malignancy.

The tumor suppressive function of TET1 and TDG is likely linked to their role in active DNA demethylation, as they had a similar impact on epigenome of murine adenomas in comparison to “wild type” Apc-Min ademonas. However, only Tet1 inactivation was associated with a CIMP-like phenotype.

Analysis of gene expression revealed transcriptional activation of the inflammatory and immune response pathways, which was confirmed by increased morphological features of inflammation in Tet1- and Tdg-mutant colonic adenomas. Previously, TET2 was shown to play a role in inflammation (32), and acute TDG knockdown (21) led to overexpression of genes belonging to inflammatory response senescence-associated secretory phenotype (SASP)(33-35). However, here we show that induction of inflammation associated with TET1/TDG inactivation is linked to CRC induction and progression.

Interestingly, the TET1/TDG inflammatory signature could separate TCGA COAD tumors based on their chromosomal instability status, which warrants further investigation. Recently, two types of aneuploidy have been described in a pan-cancer TCGA analysis: focal somatic copy number alterations (SCNA) (less than half of a chromosome arm) and arm-/chromosome-level SCNA, the latter associated with markers of immune evasion (36). It will be interesting to compare immune signature scores, and focal or arm/chromosome SCNA levels (as described in (36)) between CIN2-4 sample groups, to explore whether these types of aneuploidy preferentially segregate with hot or cold CIN, i.e. with low or high TET1 expression.

Taken together, the findings reported here suggest an important role of active DNA demethylation mediated by TET-TDG in epigenome modulation during CRC tumorigenesis: specifically, the TET-TDG axis suppresses intestinal tumorigenesis by enforcing methylation patterns and transcriptional programs that prevent activation of immune and inflammatory response pathways, and, in turn, progression to cancer.

In the future, knowledge acquired through the analysis of active DNA demethylation mediated by TET-TDG might suggest innovative cancer prevention or therapeutic strategies for inflammatory bowel disease, colitis-associated cancer and likely sporadic CRC as well. Mechanistic exploration of the processes involved in colonic inflammation and immune response regulated by TET1 and TDG has the potential to disclose novel targets to prevent transition to malignancy, implement early diagnosis and/or improve the efficacy of cancer immunotherapy.

## Acknowledgments

We thank L. Cathay for secretarial assistance. We thank the following core services at Fox Chase Cancer Center: Genomics, Cell Culture, Cell Sorting, Biological Imaging, High-Throughput Screening, Biostatistics and Bioinformatics, and Laboratory Animal Facilities. This study was supported by NIH grants CA78412 (to A. Bellacosa), CA191956 (to A. Bellacosa and T. Yen), CA218133 (to S. Grivennikov), and CA06927, and an appropriation from the Commonwealth of Pennsylvania to the Fox Chase Cancer Center. R. Tricarico was supported in part by a William J. Avery Endowed Postdoctoral Fellowship.

